# Regulation of suberin biosynthesis and Casparian strip development in the root endodermis by two plant auxins

**DOI:** 10.1101/2021.06.02.446769

**Authors:** Sam David Cook, Seisuke Kimura, Qi Wu, Rochus Benni Franke, Takehiro Kamiya, Hiroyuki Kasahara

**Affiliations:** Institute of Agriculture, Tokyo University of Agriculture and Technology, Fuchu, Japan 183-0057; Faculty of Life Sciences, Kyoto Sangyo University, Japan 603-8555; Center for Plant Sciences, Kyoto Sangyo University, Kyoto, Japan; Graduate School of Agricultural and Life Sciences, University of Tokyo, Tokyo, Japan 113-8654; Institute of Cellular and Molecular Botany, University of Bonn, Bonn, Germany 53115; Institute of Global Innovation Research, Tokyo University of Agriculture and Technology, Fuchu, Japan 183-0057

**Keywords:** Auxin, Casparian Strip (CS), Endodermis, Indole-3-acetic acid (IAA), Phenylacetic acid (PAA), Root Development, Suberin

## Abstract

The biological function of the auxin phenylacetic acid (PAA) is not well characterized in plants. Although some aspects of its biology; transport, signaling and metabolism have recently been described. Previous work on this phytohormone has suggested that PAA behaves in an identical manner to IAA (indole-3-acetic acid) in promoting plant growth, yet plants require greater concentrations of PAA to elicit the same physiological responses. Here we show that normalized PAA treatment results in the differential expression of a unique list of genes, suggesting that plants can respond differently to the two auxins. This is further explored in endodermal barrier regulation where the two auxins invoke striking differences in the deposition patterns of suberin. We further show that auxin acts antagonistically on Casparian strip (CS) formation as it can circumvent the CS transcriptional machinery to repress CS related genes. Additionally, altered suberin biosynthesis reduces endogenous levels of PAA and CS deficiency represses the biosynthesis of IAA and the levels of both auxins. These findings implicate auxin as a regulator of endodermal barrier formation and highlight a novel role for PAA in root development and differentiation.

## Introduction

The auxin phenylacetic acid (PAA) has a long history as a phytohormone, but only little is known about its biological function in plants (Cook, 2019). PAA shares several similarities with the well-studied indole-3-acetic acid (IAA) including its perception via the TIR1/AFB receptor complex (Shimizu-Mitao and Kakimoto, 2014). PAA typically has reduced potency when compared with IAA, although endogenous concentrations of PAA are often higher (Sugawara et al., 2015). PAA also possesses unique transport characteristics and is relatively immobile in plants due to its inability to be actively transported (Morris and Johnson, 1987). This auxin is most active in plant roots where exogenous treatment stimulates significant emergence of lateral roots (Sugawara et al., 2015).

Roots themselves produce a hydrophobic biopolymer that is comprised of numerous aliphatic and aromatic monomers, known as suberin (Geldner, 2013). In Arabidopsis roots, suberin is located in the endodermis where it functions as a transmembrane barrier in conjunction with the Casparian strip (Barberon, 2017). Suberin is deposited as a continuous lamella which is located on the interior of the cell wall, external to the plasma membrane (Barberon et al., 2016). This barrier works to prevent pathogen infection (Holbein et al., 2019), to regulate nutrient uptake (Barberon, 2017) and to limit radial oxygen loss (Kotula et al., 2009; Ranathunge and Schreiber, 2011). While still incomplete, the biosynthesis of suberin is well characterized in Arabidopsis and is described elsewhere (Vishwanath et al., 2015).

Plant hormones have previously been shown to influence the deposition of suberin, where the antagonistic action of abscisic acid (ABA) and ethylene control the suberization of root tissue (Barberon et al., 2016). In general, ethylene reduces suberization, while ABA promotes ectopic suberization in *Arabidopsis* roots (Barberon et al., 2016). Currently, the downstream mechanisms by which ABA and ethylene act are not understood (Barberon, 2017). In addition to these two hormones, it has also been reported that IAA treatment can upregulate expression of the putative suberin monomer transporters *ABCG6* and *ABCG20* (Yadav et al., 2014) and treatment with the synthetic auxin NAA (1-Naphthaleneacetic acid) results in altered expression of the *GDSL-lipases*, associated with the (dis)-assembly of the suberin polymer (Ursache et al., 2020). However, the effect of endogenous auxin on suberin biology has not been explored in much detail.

The Casparian strip (CS) is a cell wall modification whereby a lignin-polymer is deposited in the apoplastic space between endodermal cells in the differentiation zone of growing roots (Geldner, 2013). The deposition of CS results in a sheath that encloses root vascular bundles and fuses adjacent cell walls, creating an apoplastic barrier (Enstone et al., 2002). There is no current description of phytohormonal control over CS development aside from the recently described Casparian Strip Integrity Factor (CIF) peptide hormones, which act as a diffusible signal indicating the status of the CS (Matsubayashi et al., 2017). The CIFs are perceived by the Schengen kinases (SGN1 and SGN3), which are located on the endodermal plasma membrane interior to the CS (Doblas et al., 2017). While currently uncharacterized, the activated kinases communicate CS deficiency, likely inducing MYB36, a regulator of Casparian strip development (Kamiya et al., 2015).

Here we demonstrate that the role auxins play in barrier formation is far more complex than previously described (Yadav et al., 2014; Ursache et al., 2020). We have shown that auxin promotes suberin biosynthesis at both high and low concentrations and that PAA treatment results in a completely different suberin deposition pattern than IAA, suggesting that the two auxins likely work together to promote root suberization at different developmental stages. We also show that both IAA and PAA treatment has a negative effect on Casparian strip regulation, particularly on the *CASP* (CASPARIAN STRIP MEMBRANE DOMAIN PROTEINS) and *ESB* (ENHANCED SUBERIN 1) family genes, and on *PER64*, which are essential for normal CS deposition. However, auxin does not appear to affect CS transcription factors. In addition to influencing these root developmental processes, we have further shown that there is substantial feedback regulation on IAA biosynthesis by CS state and thus we propose a model by which auxin functions in the maintenance of the CS and of its initial establishment. Overall, these findings demonstrate that auxin is an integral regulator of endodermal barrier production.

## Results

### PAA induces a unique set of differentially expressed genes

Previous findings on the plant responsiveness to auxin have suggested that PAA is required at concentrations approximately 20 times that of IAA to induce similar levels of expression in the *GH3* (*GRETCHEN HAGEN 3*) auxin responsive genes (Sugawara et al., 2015). To determine whether PAA induces a distinct auxin response in plants we treated 2-week-old Arabidopsis seedlings with 1μM IAA or 20μM PAA and collected samples over the course of 24 hours. The resulting differentially expressed gene (DEG) list showed that 1085 and 1386 genes were significantly induced in the PAA and IAA treated plants, respectively (using a log_2_ fold change cut-off of 1.5) (Sup. Table 1). These two lists were compared to elucidate the common auxin response (821 DEGs), the PAA specific response (264 DEGs) or the IAA specific response (564 DEGs) (Figure 1A). Genes belonging to the common DEG list included typical early auxin responsive genes, such as the GH3s, the SAURs and the Aux/IAAs.

**Figure 1.**
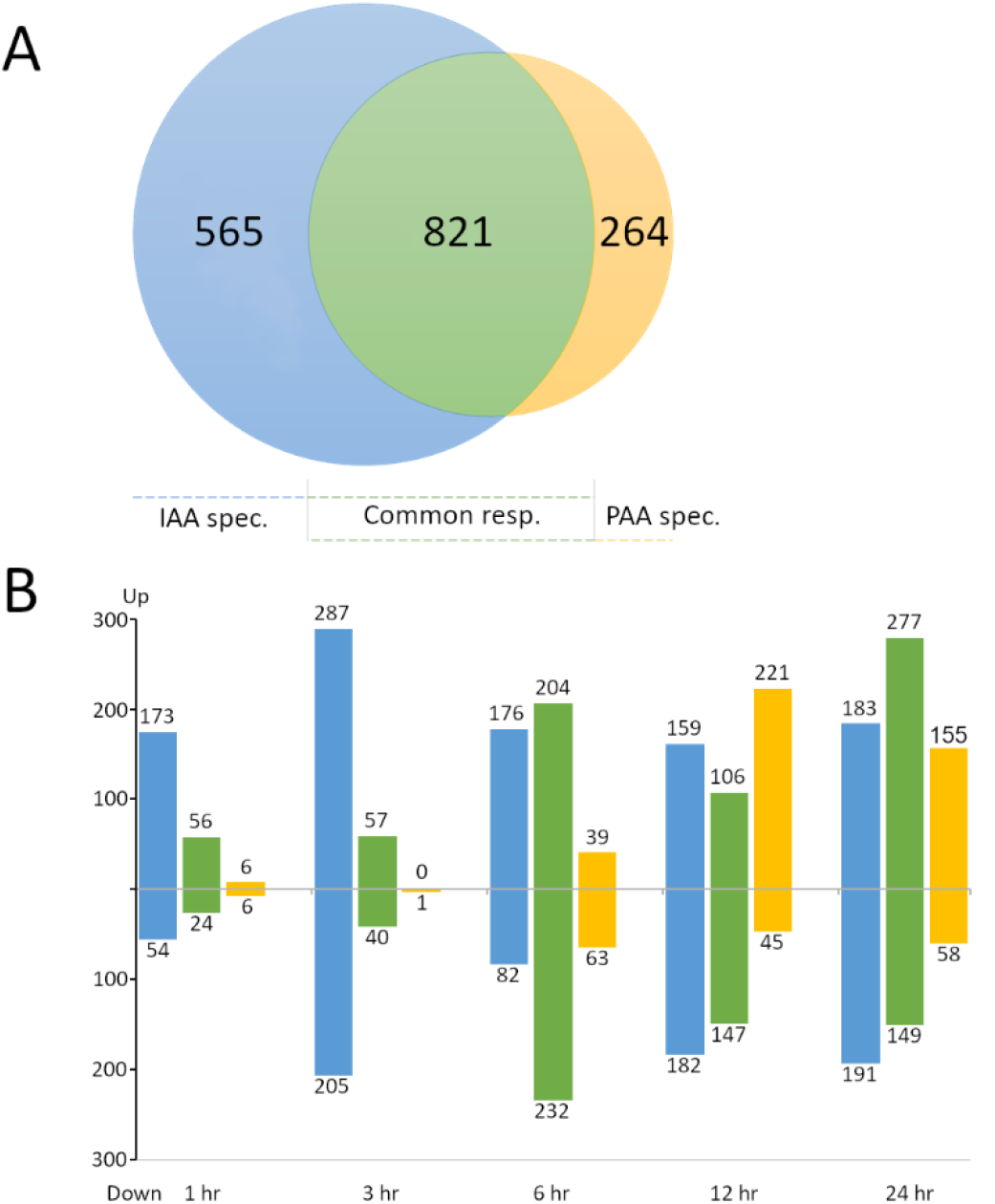
IAA and PAA treated DEGs, **A**: Venn diagram showing total number of IAA and PAA specific DEGs and common auxin response DEGs. **B**: Temporal resolution of auxin induced DEGs indicating the number of up- and down-regulated genes. DEGs defined as log_2_FC >1.5 and FDR<0.05.

Inspection of individual time points showed a strong and persistent effect of IAA treatment with the induction of 307 genes (227 specific) after 1 hour (Figure 1B). This number continued to increase to around 800 genes (374 specific) at 24 hours. The response of the plant to PAA treatment was weaker than that of IAA (based on the number of DEGs) and was delayed in its timing. PAA treatment resulted in the differential expression of less than 100 genes at the first two time points (13 specific) yet over 500 genes were induced at 6, 12 and 24 hours (Figure 1B). The number of PAA specific DEGs were also substantially increased at these later time points, in agreement with the active transport of IAA and the passive diffusion of PAA.

Given that PAA response is strongest in roots (Sugawara et al., 2015), we next investigated the expression patterns of PAA specific genes in different root cell types. We constructed heatmaps of Z-score normalized expression relative to established high resolution datasets (Brady et al., 2007) for PAA specific DEGs at each time point and mapped these to radial cell layers (Figure 2, Sup. Figure 1). The resulting heatmaps show that the DEGs induced by PAA (positive and negative) are typically expressed in root hair cells, the maturing xylem and in the phloem pole. Other clusters of DEGs localize to lateral root cells, the pericycle and the cortex.

**Figure 2.**
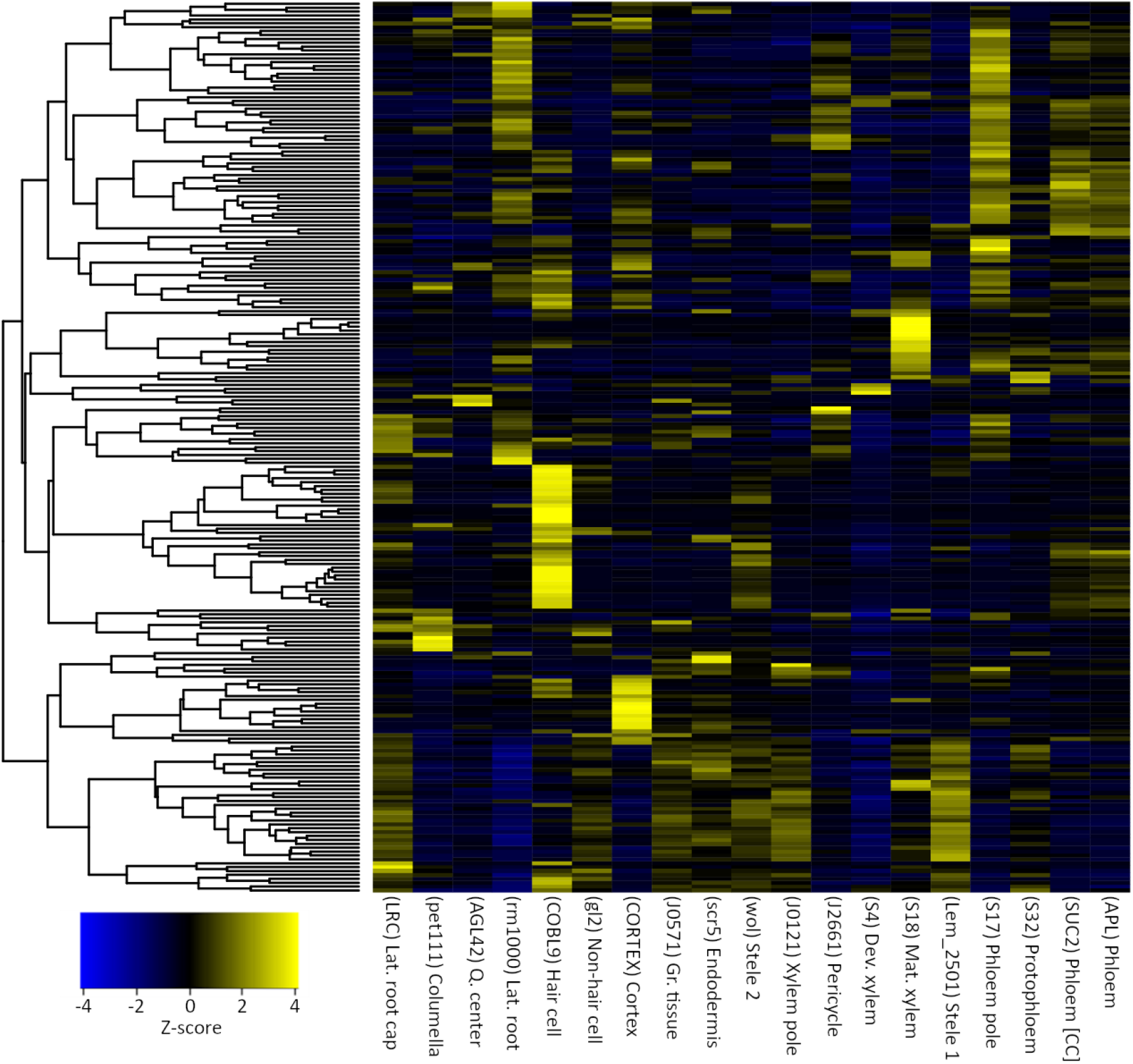
Heatmap depicting the radial distribution of PAA influenced DEGs after 12-hour treatment (Log_2_FC >1.5, FDR<0.05). Values are Z-score normalized expression relative to tissue specific indicator genes as described previously (Brady et al., 2007). See also Figure S1.

To assess the effect of PAA treatment on Arabidopsis seedlings and to determine the biological function of this hormone, we analyzed the PAA specific gene set to generate a list of enriched gene ontology terms (Sup. Table 2). The gene ontological analysis showed a strong effect of PAA treatment on suberin biosynthesis (29.44-fold enrichment), plant response to chitin (14.75-fold enrichment) and cellular response to hypoxia (13.32-fold enrichment). These terms were also highly enriched when all DEGs induced by PAA were included (Sup. Table 2B). Additionally, the term ‘cell: cell junction assembly’, which contains numerous genes required for Casparian strip development was strongly enriched in both complete PAA and IAA lists (26.04-, and 20.31-fold, respectively). None of the PAA specific terms were enriched in the IAA DEG lists (Sup. Table 2C-D).

### Auxin promotes the biosynthesis of suberin and represses the formation of the Casparian strip

More detailed investigation into the ‘suberin biosynthesis’ ontological group showed that PAA treatment upregulated numerous suberin biosynthetic genes *HORST* (*CYP86A1*), *RALPH* (*CYP86B1*), *KCS2*, *GPAT5*, *ABCG2* and *ABCG20* (Figure 3). In addition, *FAR4*, *ASFT*, *FACT*, *GPAT7*, and *ABCG6* were all elevated in both IAA and PAA treated plants. Most of these genes peaked 12 hr following auxin treatment. The suberin biosynthesis transcription factor *MYB39* (*SUBERMAN*) was also substantially elevated by both treatments. However, *MYB41*, another well-established promoter of suberin, was downregulated by both auxins at later time points. We also observed the induction of several GDSL-Lipases associated with suberin (de)polymerization (Ursache et al., 2020). These findings show that auxin, and in particular, PAA induces the expression of nearly all characterized elements of the suberin biosynthetic pathway.

**Figure 3.**
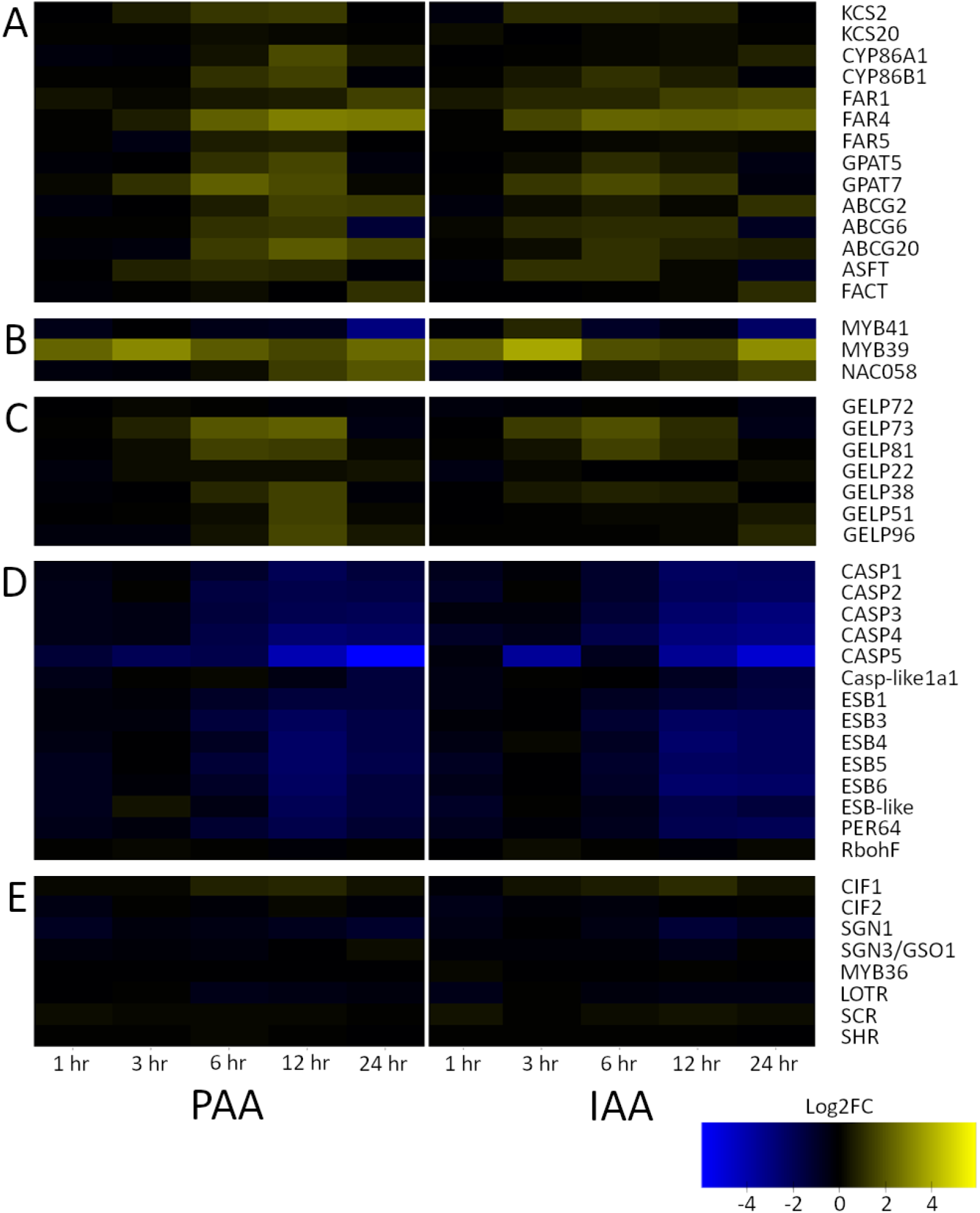
Heatmap of suberin and Casparian strip gene expression (log_2_FC) in PAA (left) and IAA (right) treated plants over a 24-hour sampling period. **A**: suberin biosynthesis genes, **B**: suberin transcription factors, **C**: putative suberin polymerization and degradation genes, **D**: Casparian strip development genes and **E**: Casparian strip affiliated transcription factors.

In addition to their effect on suberin biosynthesis, treatment with both PAA and IAA had a negative effect on the regulation of genes involved in Casparian strip development (Figure 3). All members of the Casparian strip assembly protein families which contain dirigent domains required for organization of monolignols, ESB (ENHANCED SUBERIN 1)(Hosmani et al., 2013), and CASP (CASPARIAN STRIP MEMBRANE DOMAIN PROTEIN) which are essential for Casparian strip localization and structure (Roppolo et al., 2011), were downregulated in the presence of the two auxins (Figure 3). In addition to these, the expression of the lignin biosynthetic *Peroxidase 64* (*PER64*)(Lee et al., 2013) was also reduced. The transcription factor *MYB36* which promotes the expression of the *CASPs* and *ESBs* was not differentially expressed in our dataset. Other CS associated TFs such as *LOTR* (*Lord of the Rings*), *SCR* (*SCARECROW*) and *SHR* (*SHORTROOT*) were also not significantly altered by auxin treatment. Curiously, transcripts of *CIF1* (*CASPARIAN STRIP INTEGRITY FACTOR 1*) were elevated but were just below our log_2_FC cut-off.

### Suberin is ectopically deposited in auxin treated roots

To explore the elevated expression of the suberin biosynthesis genes, we analyzed suberin deposition in *Arabidopsis* roots grown on MGRL agar plates containing 20μM PAA or 1μM IAA. Using Calcofluor White and Nile Red to stain cell walls and suberin, respectively, we observed ectopic deposition of suberin in both IAA and PAA treated roots (Figure 4). Interestingly, suberin was heavily deposited throughout the precociously emerged pericycle layer in PAA treated plants (Figure 4B, D). This is most obvious in the root tip region, which is typically devoid of suberin. However, suberin deposition in the endodermis does not appear to be altered by PAA treatment. On the other hand, treatment with IAA does not result in deposition of suberin in the pericycle but does change the deposition pattern within the endodermis (Figure 4C, F, Sup. Figure 2). In IAA treated roots, patchy suberization occurred shortly after the zone of elongation, and continuous suberization was achieved where the endodermis becomes patchy under normal conditions (ca. 30 endodermal cells). In addition to ectopic suberin deposition, the development of a mature xylem also appears to commence near the root tip in both treatments (Figure 4A–B).

**Figure 4.**
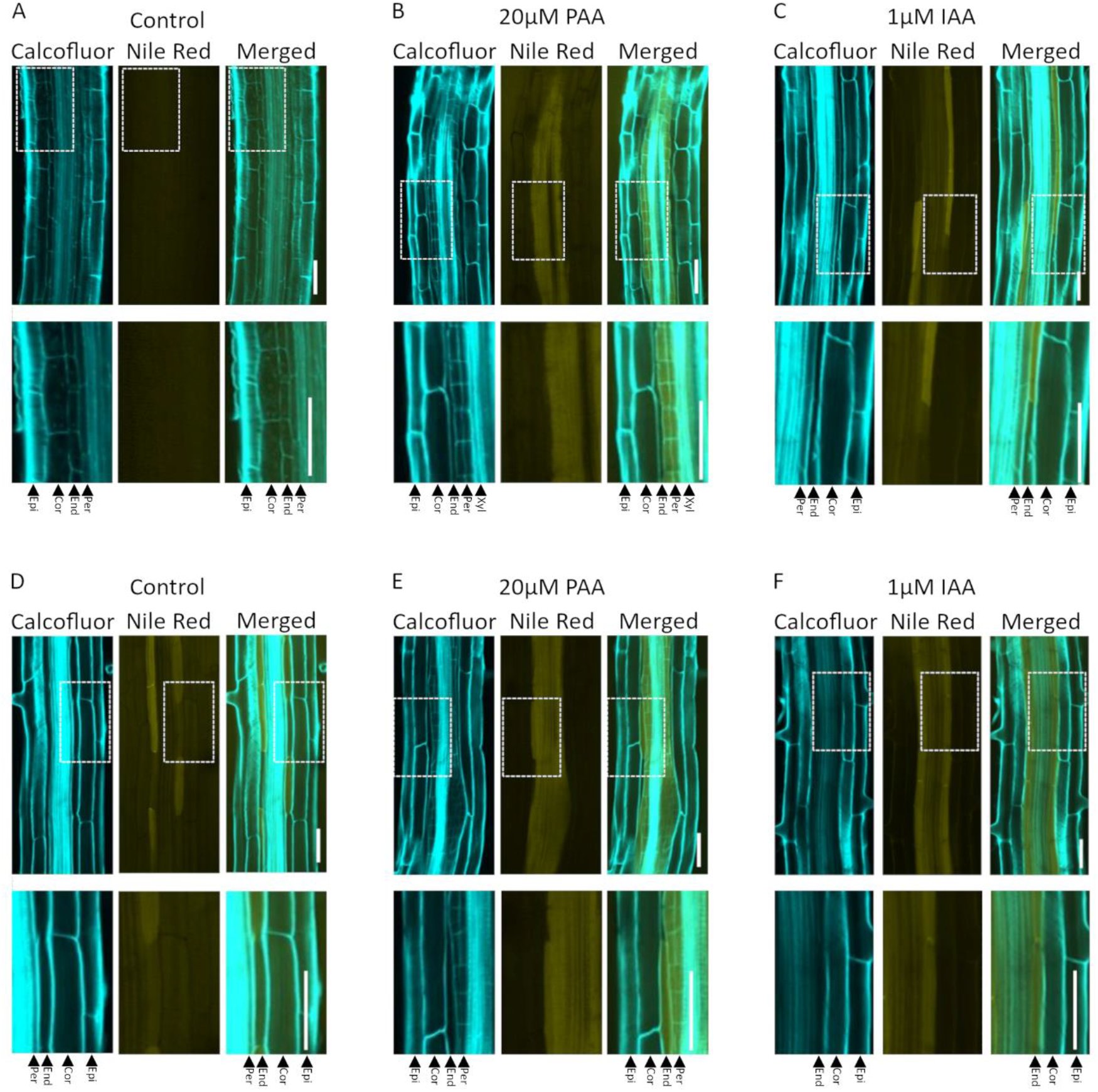
Confocal microscopy of suberin deposition in Arabidopsis seedlings grown for 5 days on MGRL media followed by 24 hours on MGRL containing a mock solution, 20μM PAA, or 1μM IAA. Panels are Calcofluor White staining of cell walls, Nile Red straining of suberin and overlay of the two. **A**: Mock root tip region, **B**: PAA treated root tip region, **C**: IAA treated root tip region, **D**: Mock ^~^40^th^ endodermal cell after the zone of elongation, **E**: PAA ^~^40^th^ endodermal cell, **F**: IAA ^~^40^th^ endodermal cell. Epi: epidermis, Cor: cortex, End: endodermis, Per: pericycle, Xyl: xylem.

Based on the auxin induced expression of suberin biosynthesis genes and the difference in DEGs between the two auxin treatments, we suspected that that the absolute levels of suberin may be elevated and that there may be some differences in the monomer composition of IAA and PAA treated roots. We thus analyzed the bound suberin monomer composition of whole Arabidopsis roots by GCMS. However, there didn’t appear to be any quantifiable difference in the overall level of suberin or in any of the individual monomers (alcohols, acids, omega-hydroxy acids (w-OH acids), Dicarboxylic acids (DCA) or in ferulic acid) (Figure 5C).

**Figure 5.**
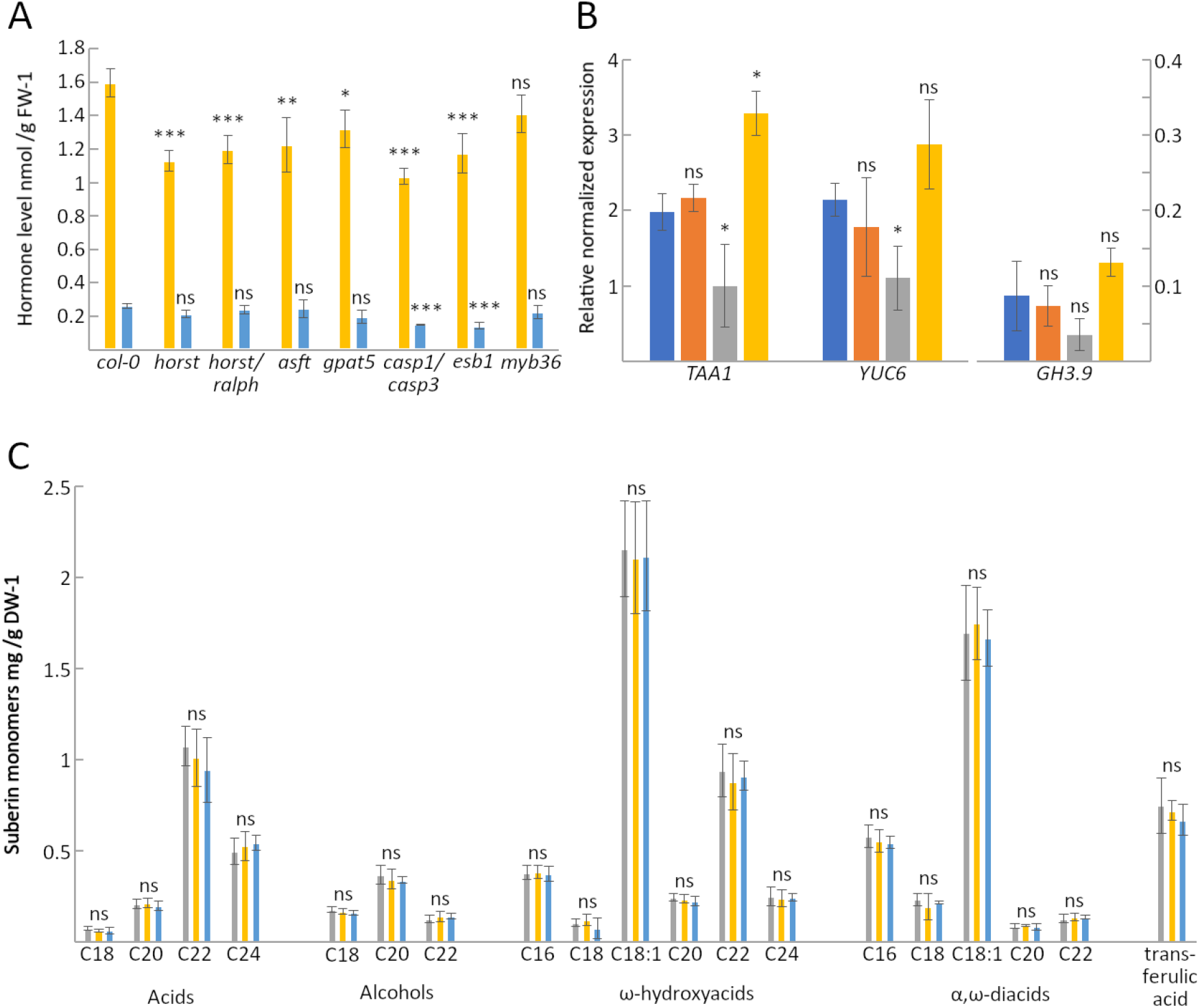
**A**: Root auxin profiles of suberin (*horst, horst/ralph, asft, gpat5*) and Casparian strip (*casp1casp3, esb1, myb36*) mutants grown on MS media. Data are means ± SD, n=4. Significances relative to col-0 (t-test, *P<0.05, **P<0.01, ***P<0.001, ^ns^ not significant). **B**: Relative normalized expression of *TAA1*, *YUC6* and *GH3.9* from the Casparian strip mutant lines in **A**. Data are means ± SD, n=4 (technical replicates=3), significances as in **A**. **C**: Suberin monomer profiles of Mock, PAA and IAA treated roots following delipidation. Data are means ± SD, n=4. Significances as in **A**, relative to mock treatment.

### Defective endodermal barriers reduce auxin levels and CS state downregulates IAA biosynthesis

We hypothesized that there may be homeostatic mechanisms that link auxin and suberin/ Casparian strip formation during root development. To explore this, we analyzed the levels of the two auxins in the suberin biosynthesis mutants; *horst*, *horst/ralph*, *gpat5* and *asft* (Figure 5A). In addition to these we also looked at the auxin levels in lines with defective Casparian strips; *casp1casp3*, *esb1* and *myb36* (Figure 5A). We suspected that if a homeostatic mechanism were in place for suberin development, suberin deficiency should result in an elevation in auxin levels, which promote suberin biosynthesis. However, the levels of PAA were significantly reduced in all suberin mutants (IAA levels were unchanged). Similar to this, *esb1* and *casp1casp3* mutants, which possess deficient Casparian strips but increased suberin levels (Baxter et al., 2009; Hosmani et al., 2013), also contained reduced levels of PAA and IAA (Figure 5A). Curiously, auxin levels were not reduced in the *myb36* mutant.

We sought to explore this relationship further by investigating the effect of CS mutants on auxin metabolism. Since the PAA biosynthetic pathway is yet to be elucidated, we analyzed the expression of the IAA biosynthesis genes *TAA1* and *YUC6,* as well as *GH3.9* (which is involved in IAA metabolism) in CS mutant roots. Consistent with the IAA levels data, we found that there was no change in the expression of *TAA1*, *YUC6* or *GH3.9* in the *myb36* line (Figure 5B). We also found expression of both *TAA1* and *YUC6* to be significantly reduced in the *esb1* mutant (Figure 5B). Intriguingly, the expression of *TAA1* was elevated in the *casp1casp3* line (Figure 5B). These results, when taken together suggest that MYB36 can control auxin levels through the downregulation of the IAA biosynthetic pathway.

To investigate this from a pharmacological perspective, we explored a transcriptome of L-Kyn treated col-0 seedlings available in the GEO database (Project: GSE32202) to determine whether auxin deficiency altered CS or suberin related transcription. L-Kynurenine (L-Kyn) is a potent inhibitor of the TAA1 family of aminotransferases, which catalyze the first step of IAA biosynthesis (He et al., 2011). Treatment with this compound results in a reduction in IAA levels, and inhibits ethylene induced IAA biosynthesis. We found that all CS genes downregulated following auxin treatment were induced in the L-Kyn treated transcriptome (Sup. Table 3). In addition to this, we also observed repression of the *CIF1* peptide hormone transcripts, which were elevated by both IAA and PAA treatment. In addition to the CS related genes, several suberin biosynthesis transcripts were similarly elevated in the L-Kyn dataset. Interestingly, the expression patterns of MYB39 and MYB41 transcription factors were reversed under low auxin conditions.

## Discussion

### Auxin induces the suberisation of the root endodermis

Suberin deposition broadly reflects the developmental stage of growing roots. Proximal to root tips, the elongation zone typically contains no suberin (Geldner, 2013). As the root matures, suberin becomes patchy and then continuously deposited in the endodermis of differentiated roots (Barberon, 2017). Our findings here show that auxin treatment dramatically alters the suberization of roots in *Arabidopsis*. Treatment with IAA results in precocious deposition of endodermal suberin in regions that are typically absent of this barrier. These changes are akin to those presented in mutants containing CS deficiencies such as *esb1* (Hosmani et al., 2013)*, casp1casp3* (Roppolo et al., 2011) or *myb36* (Kamiya et al., 2015), or in roots treated with high concentrations of ABA (Barberon et al., 2016).

Interestingly, PAA treatment does not appear to alter endodermal suberin deposition, but instead results in ectopic suberization of the root pericycle and of lateral root primordia. This has strong implications for a role of PAA in the emergence and regulation of a periderm, since it gradually supersedes the endodermis as the functional root barrier tissue (Campilho et al., 2020) and becomes extensively suberized prior to root differentiation (Wunderling et al., 2018).In line with PAA influencing the pericycle and lateral root, the periderm marker (Wunderling et al., 2018), *RAX3 (REGULATORS OF AXILLARY MERISTEM3)*, is repressed in PAA treated plants. PAA also results in the early maturation of the root xylem (as does IAA), consistent with auxin being a regulator of this process (Schuetz et al., 2013) (Figure 4B). These observations are congruent with the tissue map findings that show PAA induced genes are typically expressed in the maturing xylem, as well as the pericycle and in lateral root cells (Figure 2, Sup. Figure 1). Given that these processes normally occur later in root development (Schuetz et al., 2013; Du and Scheres, 2018; Campilho et al., 2020), it is possible that PAA functions as a driver of developmental change in *Arabidopsis* roots, no doubt dependent on concentrations of other phytohormones. The precise role of PAA in these processes requires additional investigation.

The upregulation of MYB39 and repression of MYB41 following auxin treatment and their inverse expression in the L-Kyn transcriptome clearly demonstrates a role for auxin as a biphasic promoter of suberin in *Arabidopsis* roots. Previously it has been established that ethylene and ABA are involved in the regulation of suberin biosynthesis (Barberon et al., 2016). While ABA treatment has been demonstrated to directly affect suberin transcription factors (Barberon et al., 2016; Wang et al., 2020) the exact mechanism for repression by ethylene is less clear. Suberin deposition is reduced in ACC treated plants and in the *ctr1* signaling mutant, however there is no genetic support for ethylene-based repression of suberin biosynthesis (Barberon et al., 2016). It is also well known that auxin and ethylene share a comprehensive, crosstalk network where auxin reciprocally stimulates ethylene biosynthesis (Zheng et al., 2013). Since suberin biosynthesis is upregulated under both high and low auxin conditions, it is difficult to see that elevated ethylene (which promotes auxin biosynthesis) is able to manifest such severe reductions in suberin deposition while competing with auxin induced suberin biosynthesis. Instead, it may be more likely that ethylene affects the polymerization/ degradation of the suberin lamellae.

In line with this, the bound suberin content of the auxin treated roots did not appear to be different from typical Arabidopsis root profiles, despite widespread induction of suberin biosynthesis (Figure 3) and obvious differences in the suberin deposition pattern (Figure 4). A possible explanation for this lies in the recently described GDSL-lipases, which function in the auxin-repressed polymerization and auxin-induced depolymerization of the suberin lamella (Ursache et al., 2020). Since auxin has an overall negative effect on the polymerization process, the reduced suberin level in ethylene-treated plants may well be explained by an increase in auxin levels manifesting a reduction in polymerization. Curiously, the suberin polymerization lipases; *GELP38*, *GELP51* and *GELP96*, which are repressed by NAA treatment, were upregulated following treatment by PAA (but not with IAA) (Figure 3). This inverse response to PAA may be related to the ectopic deposition of suberin in the pericycle, since Ursache *et al* (2020) focused specifically on the endodermal response. Nonetheless, PAA appears to function in a different manner to IAA once again when it comes to the regulation of suberin (de)polymerization.

Further to the specific role of PAA in suberin biosynthesis, we observed significant reductions in PAA levels in all analyzed suberin mutants, but no difference in the level of IAA (Figure 5A). The PAA biosynthesis pathway is yet to be elucidated (Cook, 2019), but suberin deficiency clearly influences PAA homeostasis. Given that PAA is not actively transported in plants (Morris and Johnson, 1987), we suggest that PAA levels are likely reduced due to repressed biosynthesis, since an increase in PAA metabolism (at least via the GH3s), should result in a concerted reduction in IAA levels (Sugawara et al., 2015; Aoi et al., 2019). The mechanism behind this feedback is currently unknown, although recently MYB36 has recently been linked to suberin biosynthesis (Wang et al., 2020) and the regulation of the entire endodermal barrier may be controlled by a single system transcription factor.

### Auxin is an antagonist of Casparian strip gene expression

The regulation of the Casparian strip (CS) is controlled by the SHR:SCR:MYB36 transcription factor cascade (Li et al., 2018). SHR as an integral root developmental transcription factor controls the expression of MYB36 and SCR in parallel (Li et al., 2018). MYB36 then regulates the transcription of the *CASP* and *ESB* gene families and of several peroxidases to reenforce CS formation (Lee et al., 2013; Kamiya et al., 2015; Liberman et al., 2015) while SCR controls the expression of the Schengen kinases, *SGN1* and *SGN3*, to direct the subcellular localization of the CS (Li et al., 2018).

Our data show that auxin treatment results in massive reductions in the expression levels of the CASP and ESB gene families (Figure 3), similar to those reported in the *myb36* mutant (Kamiya et al., 2015; Liberman et al., 2015). However, the expression of *MYB36* and other CS regulating transcription factors were not affected by IAA or PAA treatment, indicating that auxin is directly repressing the expression of CASP and ESB genes, or operating through an as yet unknown factor. In the L-Kyn transcriptome we found that all CS genes downregulated following auxin treatment were instead elevated when auxin biosynthesis is inhibited, which rather suggests that auxin interferes with MYB36 signaling (Sup. Table 3). In addition to this, we observed repression of the *CIF1* peptide hormone transcripts in the L-Kyn dataset, which was slightly elevated following auxin treatment. The combination of these findings implicates auxin as a clear antagonist of CS development; however, auxin also appears to control the abundance of the CIF1 peptide hormone, which may increase the sensitivity of the plant endodermis to CS deficiency.

We also observed a clear reduction in auxin levels in the CS mutants *esb1* and *casp1casp3* but not in the *myb36* signaling mutant. In agreement with this, we observed a reduction in the *TAA1* and *YUC6* transcripts in the *esb1* line, but not in *myb36* (Figure 5B), suggesting that a functional MYB36 is necessary for the repression of auxin biosynthesis in CS deficient plants. It is unclear why TAA1 transcripts were elevated in *casp1casp3,* but other IAA biosynthesis genes or those of ethylene turnover may also be involved. To determine whether MYB36 can control auxin in general, we explored *TAR* and *YUC* gene family expression in the recently published *iMYB36* and *iSHR* overexpression transcriptomes (Li et al., 2018; Wang et al., 2020). Even though endodermal CS is complete in these lines, the IAA biosynthesis genes, *TAR2* and *YUC3*, were repressed in *iSHR*, but not in *iMYB36* (Sup. Table 4). This suggests that while MYB36 is essential for the repression of IAA biosynthesis, MYB36 alone does not appear to be sufficient to alter auxin levels. On the other hand, SHR, as a major root developmental transcription factor is able to control auxin levels through recruitment of SCR, since SCR can repress both Trp and IAA biosynthesis (Moubayidin et al., 2013) as well as promoting auxin efflux through repression of ARR1 and thus SHY2/IAA3 (Dello Ioio et al., 2008; Moubayidin et al., 2016). However, neither *SHR* nor *SCR* are induced in the *iMYB36* line, which points instead to an as-yet-unknown mechanism for controlling auxin biosynthesis in response to a deficient CS.

## Conclusions

Here we have presented evidence that auxin is significantly involved in the regulation of both suberin and the Casparian strip (Figure 6). Despite distinct suberin deposition patterns, both auxins appear to have an inductive effect on suberin though the biphasic regulation of the *MYB39* and *MYB41* transcription factors. The auxin phenylacetic acid also appears to manifest different responses in suberin polymerization and appears to be under a homeostatic mechanism that does not apply to IAA. The precocious emergence of suberized cell layers in the pericycle and early xylem maturation seen in PAA treated plants invites many questions around the importance of this relatively unknown phytohormone in root development. Future work on the regulation of suberin should incorporate both auxins, as well as ethylene and ABA to determine the exact mechanisms by which suberin condition is communicated and manipulated in the endodermis and the emerging pericycle.

**Figure 6.**
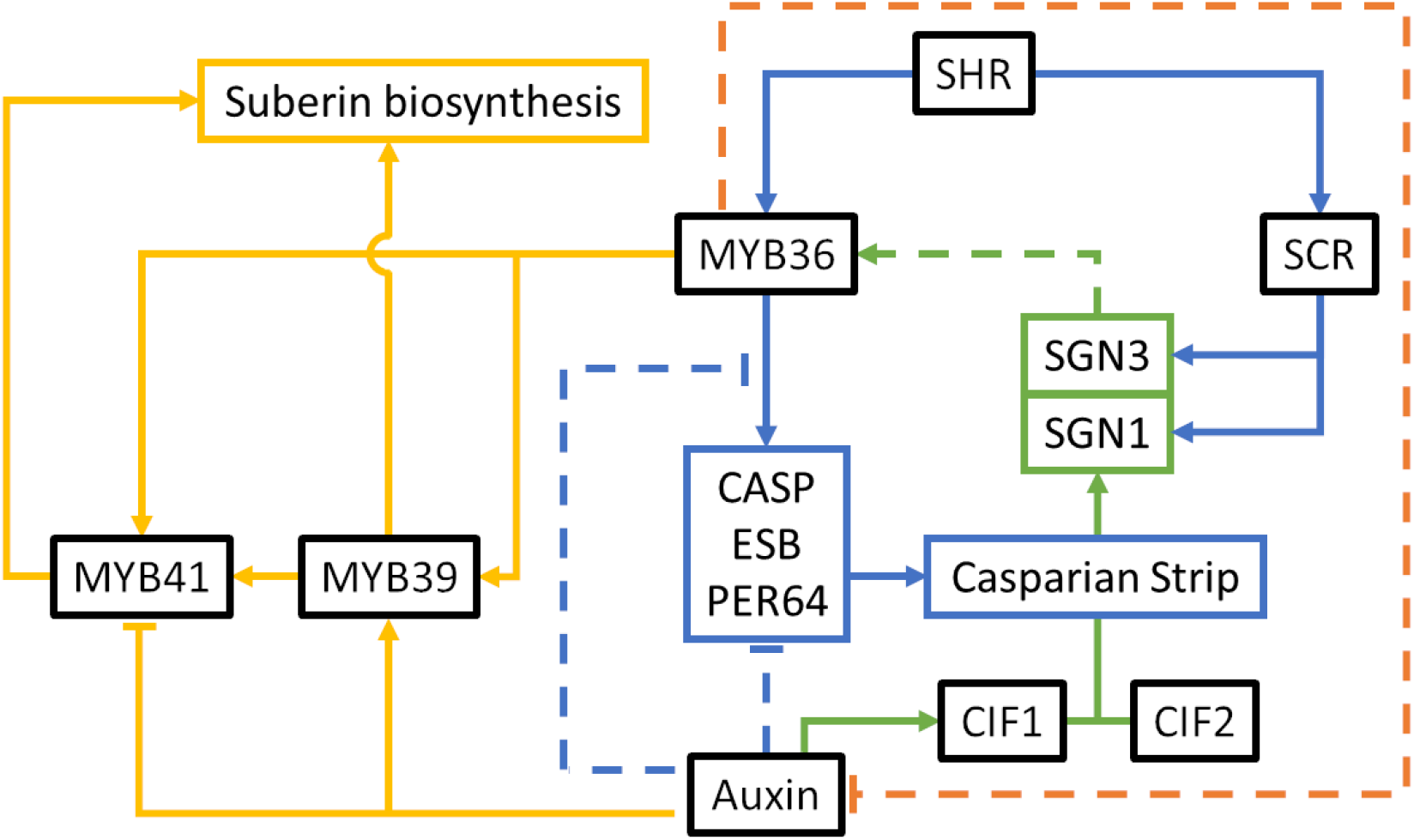
Hypothetical model integrating auxin and endodermal barrier regulation. Colors describe subsections of the regulatory network; Blue: Casparian strip development/ root development, Yellow: suberin biosynthesis, Green: CIF peptide hormone pathway. Orange: uncharacterized CS/ auxin feedback loop. Arrows and flat ends indicate direction of influence (promotion or repression, respectively) and dashed lines remain to be characterized. CASP: Casparian strip membrane protein, CIF1/2: Casparian strip integrity factor, ESB: enhanced suberin, MYB36/39/41: MYB type transcription factor, PER64: peroxidase64, SCR: scarecrow, SHR: shortroot, SGN1/3: Schengen receptor kinases1/3.

Furthermore, our data indicate that CS development is also regulated by auxin. Both IAA and PAA treatment reduces the transcription of the *ESBs*, *CASPs* and of *PER64*. The induction of these genes in L-Kyn treated plants further suggests that auxin may be interfering with MYB36 signaling, However, additional work is required to elucidate this dynamic. The effect of compromised CS integrity (as in *myb36*, *esb1* and *casp1casp3*) on auxin levels also warrants further investigation, although given the role that auxin: cytokinin interplay has on the transition from division to differentiation in the growing root, it is tempting to speculate that auxin is the direct mechanism by which endodermal cells gain their identity. In the quiescent center, SCR represses the expression of *ARR1* which promotes *PIN* expression and auxin efflux (Moubayidin et al., 2016). Auxin is transported to the transition zone (TZ) where it induces *ARR1* in pre-differentiated tissue (Moubayidin et al., 2013). ARR1 then promotes the expression of the cytokinin inducible *GH3.17* (Di Mambro et al., 2017) and of *SHY2/IAA3* which restricts auxin efflux (Tian et al., 2002; Dello Ioio et al., 2008) causing an auxin minimum at the TZ (Di Mambro et al., 2017). This auxin minimum alleviates the repression of MYB36 signaling, promoting CS development. In agreement with this, the CASP and ESB proteins tend to accumulate at the plasma membrane of cells in the TZ (Roppolo et al., 2011; Hosmani et al., 2013). As the formation of a CS is a key indicator of endodermal cell identity, auxin appears to be directly involved in the differentiation of this cell type.

On a final note, given that the two auxins are perceived by the same signaling machinery (TIR1/AFBs)(Shimizu-Mitao and Kakimoto, 2014), yet can induce unique transcriptional and physiological changes, it is possible that an IAA:PAA ratio, an ‘**auxin function**’ is what governs these differential plant responses. While this suggestion requires much experimental and conceptual work, it is no longer acceptable to ignore the contribution of PAA in the plant auxin response.

## Materials and Methods

### Plant material and Growth conditions

For RNA-Seq, qPCR and GC-MS experiments, wild type *Arabidopsis thaliana* seedlings (col-0) were stratified in the dark at 4°C for 2 days. Seedlings were grown on MS medium (1% sucrose, 0.8% agar) for 10 days at 23°C under a 16-hour photoperiod. Seedlings were transferred to culture flasks containing liquid MS (30 mL, 1% sucrose) and incubated under the same conditions for an additional 2 days for equilibration. Seedlings were then treated with 30μL solutions containing 1mM IAA (in DMSO), 20mM PAA, or a control (Final concentrations of 1μM and 20μM respectively) and returned to the above conditions. Samples were harvested at intervals of 0, 1, 3, 6, 12 and 24 hours, weighed and frozen in liquid N2. For Nile Red staining experiments, seeds were sterilised and plated on MGRL media in the dark at 4°C for 2 days and then transferred to growth chambers as above in a vertical orientation. Seedlings were then transferred to MGRL media containing 1μM IAA, 20μM PAA or a mock solution of DMSO for 24 hours. For hormone analysis, mutant seeds of *horst*, *horst/ralph*, *asft*, *gpat5*, *casp1casp3*, *esb1* and *myb36* were stratified in the dark at 4°C for 2 days and transferred to 1x MS plates (1% sucrose, 0.8% agar). Plates were transferred to growth chambers as above and grown for 12 days.

### RNA sequencing, DEG determination and GO analysis

Total RNA was extracted using an RNeasy kit (QIAGEN) and RNA was stored at −80°C until RNA Sequencing. Each RNA sample was prepared from a pool of five whole seedlings. Three independent RNA samples for each condition were used for the analyses. After RNA integrity was confirmed by running the RNA samples on Agilent RNA 6000 Nano Chip (Agilent Technologies), 0.5 μg of the samples were used for library preparation using Illumina TruSeq Stranded mRNA LT Sample kit according to the manufacturer’s protocol. Sequencing was performed on a NextSeq 500 (Illumina), and 75 bp-long single-end reads were obtained. The reads were mapped to the reference *A. thaliana* genome (TAIR10) using TopHat2 (Kim et al., 2013) and counted using the htseq-counts script in the HTSeq library (Anders et al., 2015). Count data were subjected to trimmed mean of M-values normalization, and differentially expressed genes were defined using EdgeR (Robinson et al., 2009) and ‘limma’ (Ritchie et al., 2015). Genes with a log_2_FC of >1.5 and a FDR < 0.05 were classified as differentially expressed genes (DEGs). Subsequent DEG lists were parsed through the panther gene ontological (GO) classification system to obtain over/ underrepresented GO terms.

### Histochemical staining and confocal microscopy

Samples were fixed with 4% paraformaldehyde in 1x PBS and were cleared using the ClearSee protocol (Ursache et al., 2018). Samples were stained in 0.05% Nile Red in ClearSee solution for 12-16 hours and were washed in ClearSee. Stained roots were then transferred to an 0.1% solution of Calcofluor White in ClearSee for 30 minutes followed by 30 minutes wash in ClearSee. Confocal laser scanning microscopy experiments were performed by using FV1000 (Olympus) confocal microscope. For suberin and cell wall visualization, the excitation and emission wavelengths were set as follows: 405 nm excitation and 425-475 nm emission for Calcofluor White, 561 nm excitation and 600-650 nm emission for Nile Red.

### GC-FID and GC-MS analysis

Samples were harvested after 12 hours of treatment in liquid culture (as above) and were washed thrice in dH20 and dried to remove excess media. Roots were excised and cut into small (<10mm) pieces after which ^~^200mg of tissue was transferred into glass vials containing 10mL 1:2 solution of chloroform: methanol for 24 hours with agitation and intermittent refreshing of the solvent solution. The samples were then placed in 10mL 2:1 chloroform: methanol solution for 24 hours with agitation and intermittent refreshing of the solution. Delipidated samples were then dried completely and stored prior to analysis by GC-FID and GC-MS as described previously (Franke et al., 2005).

### Hormone extraction and LC-MS analysis

12 day old roots were harvested from mutant seedlings into 4 replicates of ~50 mg fresh weight and were immediately frozen in liquid N2. Samples were processed as described previously (Aoi et al., 2019). Purified samples were injected into an Agilent 6420 Triple Quad system (Agilent Technologies, Inc.) with a ZORBAX Eclipse XDB-C18 column (1.8mm, 2.1mm x 50mm). The HPLC separation and MS/MS analysis conditions are also described previously (Aoi et al., 2019).

## Supplemental material

**S. Figure 1.** Heatmaps depicting the radial distribution of PAA induced and repressed DEGs after 1(A,B), 3(C), 6(D,E), 12(F,G) and 24(H,I)-hours treatment (Log_2_FC >1.5, FDR<0.05). Values are Z-score normalized expression relative to tissue specific indicator genes as described previously (Brady et al., 2007). See also Figure 2.

**S. Figure 2.** Suberin deposition pattern in Arabidopsis seedlings grown for 5 days on MGRL media followed by 24 hours on MGRL containing 1μM IAA. Data are mean endodermal counts ± SD, n=6.

**S. Figure 3.** Confocal microscopy of suberin deposition in Arabidopsis seedlings grown for 5 days on MGRL media followed by 24 hours on MGRL containing mock solution, 20μM PAA or 1μM IAA. Panels are Calcofluor staining of cell walls, Nile Red straining of suberin and overlay of the two. A: Continuous suberization of the region around the 70^th^ endodermal cell after the zone of elongation under mock treatment, B: continuous suberization after PAA treatment and C: continuous suberization after IAA treatment at similar positions. Epi: epidermis, Cor: cortex, End: endodermis.

**S. Table 1.** Cumulative list of IAA and PAA DEGs from all time points.

**S. Table 2.** Gene ontological analysis of PAA and IAA DEGs. A: PAA specific, B: PAA all, C: IAA specific, D: IAA all.

**S. Table 3.** LogFC of suberin and Casparian strip related genes in *tar2/wei8* and L-Kyn treated plants compared to 12 hour treatment with 1μM IAA and 20μM PAA. *tar2/wei8* and L-Kyn data were obtained from the GEO database (Project: GSE32202).

**S. Table 4.** LogFC of auxin biosynthesis and metabolic genes in the iMYB and iSHR transcriptomes published previously (Li et al., 2018; Wang et al., 2020).

## Data Availability

RNA-seq data reported in this paper have been deposited in the DDBJ Sequence Read Archive (DRA) under the project ID [DRA011584]

## Acknowledgments

The authors would like to thank Sakamoto Tomoaki for his assistance with the submission of the RNASeq read data to the DDBJ Sequence Read Archive (DRA).

